# Statistical Atlas-Based Surrogate Model of Biventricular Wall Mechanics

**DOI:** 10.64898/2026.01.07.697811

**Authors:** Alejandra Robles, Raksha Konanur, Anna Qi, Henrik Finsberg, Joakim Sundnes, Andrew D. McCulloch

**Author notes:** These authors contributed equally to this work (Alejandra Robles, Raksha Konanur).

## Abstract

Here we use a statistical atlas of end-diastolic (ED) and end-systolic (ES) biventricular shapes – previously derived from the UK Biobank imaging substudy – to generate meshes for finite element (FE) simulations of ventricular wall mechanics. The models used the Holzapfel–Ogden constitutive law for passive material properties and a time-varying elastance model of systolic tension development. Simulated ED and ES deformations were projected onto the shape atlas and the principal components were used to train a multi-layer perceptron as a surrogate model. The input layer included shape modes of the unloaded ventricular geometry, and material parameters and ventricular pressures at ED and ES. After training with 444 simulations, the surrogate model achieved a mean square error in predicted displacements of *<* 2 mm and volumetric overlaps with FE-predicted deformed shapes *>* 97%, demonstrating good fidelity to the simulated ground truth. This approach may enable accurate prediction of ventricular wall mechanics without computationally expensive finite element analysis, offering a more feasible method for rapid, subject-specific cardiac modeling.

## 1 Introduction

Passive and active myocardial material properties are key determinants of regional ventricular wall mechanics and global pump function. Changes in material properties are important markers of cardiovascular aging, disease risk and progression [1], [2], [3]. However, because these properties cannot be measured directly, healthcare workers have used inverse models of ventricular biomechanics to estimate them.

Advances in medical imaging and inverse finite element (FE) modeling enable the construction of subject-specific models (“digital twins”) of ventricular biomechanics. These advances allow us to estimate quantities that cannot be measured clinically including regional wall stress, strain distributions, and mechanical properties. Cardiac magnetic resonance imaging (MRI) and computed tomography resolve cardiac geometry throughout the cardiac cycle, including at end-diastolic (ED) and end-systolic (ES) states. Automated pipelines now allow these image volumes to be segmented and meshed into FE models automatically. The models can then simulate ventricular wall mechanics in these geometries given material properties and hemodynamic boundary conditions. Constitutive models have also been derived from in-vitro measurements in cardiac muscle. However, identifying patient-specific parameters typically requires costly inverse optimizations, which are performed by minimizing the residuals between model-predicted and observed wall displacements [1], [4], [5].

Material parameter optimization of FE models typically requires numerous iterations of large, nonlinear, 3D forward FE simulations. We aim to overcome this limitation by using deep learning to train a network that can serve as a surrogate FE model. However, since typical cardiac FE models can require tens of thousands of degrees of freedom to model ventricular shape and wall motion, the computations needed to train such a surrogate model could be computationally prohibitive.

Here we take advantage of statistical atlases of biventricular shape and wall motion derived using principal component analysis (PCA) from large databases of cardiac MRI exams, such as the UK Biobank Imaging Substudy [6], [7], [8]. These atlases greatly reduce dimensionality while preserving the features that explain variation across the population. The atlas can also be used to generate statistically representative synthetic geometries that can be shared publicly without the need for additional protections of individual health data. The atlas used here was derived from 38,858 UK Biobank cardiovascular MRI studies in healthy and diseased volunteers that includes both ED and ES biventricular shapes. The first 25 shape modes explained more than 90% of the shape variance across the cohort [8].

In this paper, we use finite element meshes generated from the UK Biobank biventricular atlas to generate synthetic, statistically representative data linking geometry, pressure, and material parameters. With the atlas as a latent space, FE solutions resolved into shape modes were used to train and validate a surrogate model for predicting deformed deformed shapes of the heart given an unloaded geometry, chamber pressures, and material properties. Even with a modestly sized training set, the surrogate model predicted FE-computed deformed geometries with average errors under 1 mm and maximum errors less than 2 mm.

## 2 Methods

We developed a computational pipeline to simulate biventricular wall mechanics using a population-derived virtual cohort. Statistically representative geometries were obtained from pseudo-random samples of the first 25 modes of ED/ES shapes in a biventricular shape atlas derived from imaging substudy volunteers in UK Biobank [8]. Because FE models require an unloaded stress-free reference geometry, sampled ED shapes were adjusted with a heuristic to match extrapolated unloaded ventricular volumes. These unloaded geometries were then generated and meshed, assigned myocardial fiber architecture, and used for FE simulations of passive diastolic filling and active systolic contraction for physiologically relevant ranges of ventricular ED and ES pressures. Next, simulated deformed geometries were mapped back into atlas space, and used to train surrogate machine learning models. The full workflow is shown in Fig. 1.

**Figure 1:**
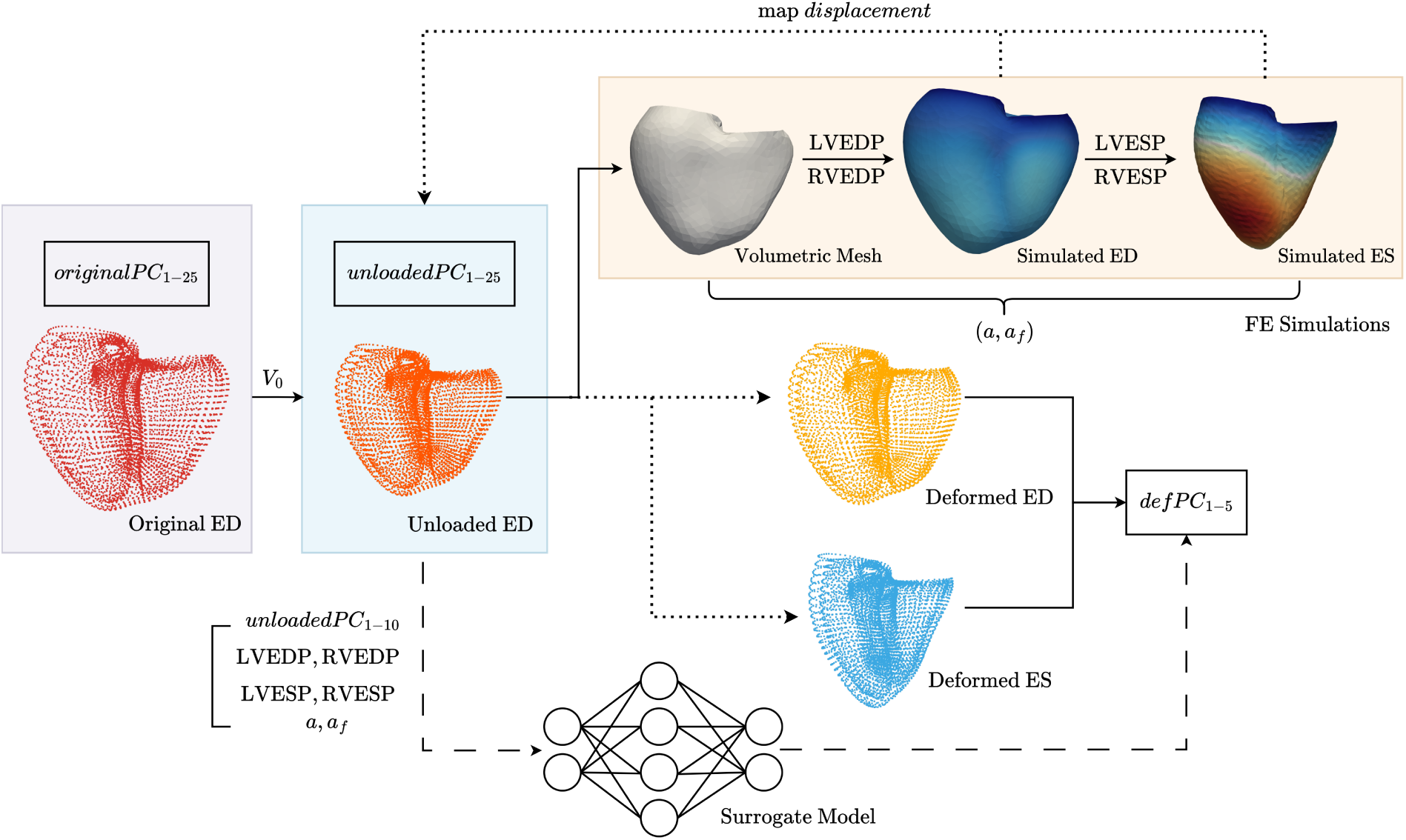
Complete workflow beginning with the original PC scores from sampled atlas data to determining the deformed PC scores, then training and validating the surrogate model. Randomly sampled shape modes (*originalPC*_1*−*25_) are adjusted to *unloadedPC*_1*−*25_ via a heuristic to reduce ventricular size so that LVEDV approximates estimates unloaded volume *V*_0_. Using this unloaded ED geometry, a volumetric FE mesh is constructed and deformed to ED given prescribed filling pressures (LVEDP, RVEDP) and to ES with prescribed systolic pressures (LVESP, RVESP) for ranges of passive and active stiffness scaling coefficients. Simulated ED and ES deformed geometry pairs are projected on to the atlas, resulting in the *def PC*_1*−*5_. A MLP surrogate model was trained with inputs *unloadedPC*_1*−*10_, pressures, and material properties and outputs *def PC*_1*−*5_.

### 2.1 Geometry and Hemodynamic Sampling

The biventricular meshes were generated from sampling the UK Biobank [9] and extracting the distribution of chamber volumes. Then using the first 25 shape modes of a biventricular statistical atlas of ED and ES shapes [6], [8], a synthetic cohort with PC scores representative of the aforementioned distribution was generated. A single set of PC scores, when projected back to the atlas, results in point clouds corresponding to RV and LV epicardial and endocardial surfaces at both ED and ES. Using arterial systolic pressures from the UKB cohort, plausible systolic blood pressures were sampled and used as LV ES pressures. LV and RV ED pressures were selected from normal values in human hearts [10]. Finally right ventricular ES pressures were computed by assuming a 9:4 ratio between the LV and RV respectively. This ratio was chosen based on the average ED pressure in the LV and RV. This ensured that the generated simulated hearts had LV to RV chamber volumes consistent with experimental data.

### 2.2 Generation of Unloaded Biventricular Meshes

To estimate the unloaded shape, we first determined the unloaded volume, *V*_0_, of the LV. For the UK Biobank participants, *V*_0_ was obtained by fitting the Klotz curve [11] to LVEDP and LVEDV. We then established a linear relationship (1) between *V*_0_ and LVEDV with an *r*^2^ = 0.9968.

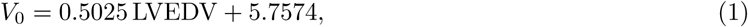

Using this estimated unloaded volume *V*_0_, we used iterative gradient descent to find modified values of the shape mode scores of the model that reduced ED LV chamber volume to *V*_0_ ±0.1 mL. LV volumes were computed from shape PCs using the Biventricular Modeling Pipeline (biv-me) [12]. The PC scores at the end of the optimization were assumed to correspond to subject-specific unloaded geometry at zero pressure. The modified shape modes were projected back to the atlas and used to generate ED surface point clouds assumed to correspond to an equivalent unloaded geometry making use of the the ukb-atlas repository. Then, using meshio [13], surface meshes corresponding to the aortic valve (AV), mitral valve (MV), pulmonary valve (PV), tricuspid valve (TV), LV, RV, RV free wall, and the epicardium were constructed [14]. With gmsh [15], an unstructured volumetric mesh was constructed from these surfaces with a fixed characteristic length of 5.0 mm.

Fiber architecture was assigned to the unloaded meshes using the Laplace Dirichlet Rule-Based algorithm [16]. The myofiber helix angle was assumed to vary from +60*^◦^* at the endocardium to −60*^◦^* at the epicardium on both the LV and RV.

### 2.3 Myocardial Mechanics

#### Passive Properties

To describe the passive material properties of the myocardium, we used a transversely isotropic version of the hyperelastic strain energy function proposed by Holzapfel and Ogden (2009) [17]:

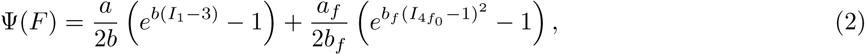

where *a, a_f_, b, b_f_* are material parameters and the invariants *I*_1_ and *I*_4*f*0_ are defined as

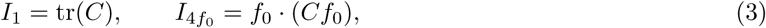

with *C* denoting the right Cauchy–Green tensor and *f*_0_ the myocardial fiber direction. The myocardium is assumed to be incompressible, which we enforce with a Lagrange multiplier representing the hydrostatic pressure *p* in an augmented Lagrangian formulation [18], [19].

Constants *b* and *b_f_* were set to 9.726 and 15.779 respectively based on biaxial testing data [20]. The parameters *a* and *a_f_*, scaling the isotropic stress and the fiber stress, were varied as follows to generate six parameter combinations for each simulation: *a* = 0.50 and 1.28 kPa and *a_f_* = 1.7, 10.0 and 20.0 kPa. These values were based on previous estimates for healthy myocardium [21] and biaxial testing [20].

#### Active Properties

Active contraction of the myocardium was represented using the formulation by Nash and Hunter (2000) [22]:

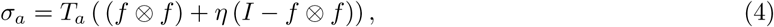

where *T_a_* is the magnitude of the active fiber stress and *η* controls the proportion of active fiber tension generated in the transverse directions. Following previous studies, we assumed the transverse isotropy and fixed *η* to 0.2. *T_a_* was set to 0.0 during the initial loading phase from unloaded to ED, and ramped up linearly 0.0 at ED to 200.0 kPa at ES.

### 2.4 FE Implementation

Simulations were performed in FEniCSx [19] with toolboxes for cardiac geometries and atlas meshing [14], [23]. The nonlinear solver uses Newton iterations with appropriate stabilization and convergence criteria. Taylor-Hood elements were used for pressures and displacements, with first-order Lagrange elements used for pressure and second-order Largrange elements for displacements.

#### Boundary Conditions

All valves (AV, MV, PV, TV) were assigned fixed Dirichlet boundary conditions, with the displacements set to zero. Neumann boundary conditions were used to impose pressures on the left and right ventricles, similar to a previous study [24].

#### Loading

Loading was divided into two phases: passive loading of the unloaded geometry to ED pressure with no active stress, and the active loading of the ED geometry to ES with a linearly ramped active stress and pressure. The prescribed ED and ES pressures applied are summarized in Table 1.

**Table 1:** Pressure and active tension cases.

| Case | Unloaded → ED (kPa) | | | ED → ES (kPa) | | |
| --- | --- | --- | --- | --- | --- | --- |
| | $LV_{EDP}$ | $RV_{EDP}$ | $T_a$ | $LV_{ESP}$ | $RV_{ESP}$ | $T_a$ |
| LOW | 0.80 | 0.356 | 0.0 | 14.0 | 2.0 | 200.0 |
| NORMAL | 1.20 | 0.533 | 0.0 | 18.0 | 3.0 | 200.0 |
| HIGH | 1.60 | 0.711 | 0.0 | 25.0 | 4.0 | 200.0 |

Three sets of ED pressure were simulated: low, medium, and high. Applied LVESP were based on the distribution of systolic blood pressures in the UK Biobank cohort. The meshes were deformed from unloaded to ED by linearly load-stepping from unloaded to ED. No active stress was applied during the filling phase. For each ED load, three sets of ES pressure pressures were applied to simulate systole corresponding to low, medium, and high ES pressures. In the ejecting phase, an active tension (*T_a_*) (refer Eq. 4) of 200.0 kPa was applied by linearly loadstepping from ED to the lowest ES case. While maintaining constant *T_a_*, ES pressures were further increased to medium and then high cases. The final displacements at each pressure case for both ED and ES were saved. Thus, each combination of material parameters for each geometry resulted in 9 different ED-ES combinations.

### 2.5 Mapping to Atlas Space

To recover the PC scores from the deformed FE meshes, the deformed coordinates must be of expected length and order. This is required as the atlas expects coordinates in a particular order to maintain connectivity. However, since the simulations were performed on unstructured volumetric meshes, neither the order nor the connectivity is maintained. The mapping between the original coordinates and the nodes in the volumetric meshes is also unclear. To solve this problem, ED and ES displacements were mapped using a distance-weighted interpolation of the nearest 4 unloaded coordinates back to the original ED point cloud derived from the atlas and used to generate the unloaded mesh. These interpolated displacements were added to the coordinates of the point cloud corresponding to the unloaded atlas PCs. This preserves the ordering and number of the original coordinates, which can then be projected back onto the atlas to recover the PC scores for simulated deformed ED and ES pairs. Chamber volumes corresponding to the solutions were extracted from these point clouds using the biv-me package [12].

### 2.6 Surrogate Model

We trained a feedforward neural network to predict deformed principal component (*def PC*) scores from unloaded shape scores, ventricular pressures and material parameters.

#### Data preparation

Fifty sampled geometries were separated into 40 for training and 10 for testing. The input layer consisted of the first 10 shape modes (*unloadedPC*_1*−*10_) of the unloaded geometry, LV and RV pressures at end-diastole and end-systole (LVEDP, LVESP, RVEDP, RVESP), and two material parameters (*a*, *a_f_*), resulting in a 16-dimensional input vector. The active stress was not included in the training data as it was constant both at ED and at ES across all simulations. The target outputs were the first five principal components of the defomed meshes (*def PC*_1*−*5_).

#### Model architecture

The machine learning model was a multilayer perceptron (MLP) with two hidden layers of 128 and 64 units, implemented using Pytorch [25]. Each hidden layer used a ReLU activation followed by dropout regularization (*p* = 0.1). The output layer was a fully connected linear mapping with 5 units (corresponding to *def PC*_1*−*5_). In total, the network contained 10,757 trainable parameters. The architecture is summarized in Table 2.

**Table 2:** Architecture of the MLP used for defPC prediction.

| Layer | Input size | Output size | Parameters |
| --- | --- | --- | --- |
| Linear + ReLU + Dropout | 16 | 128 | 2,176 |
| Linear + ReLU + Dropout | 128 | 64 | 8,256 |
| Linear (output) | 64 | 5 | 325 |
| **Total** | – | – | **10,757** |

#### Training

The network was trained to minimize the mean squared error (MSE) between predicted and simulated deformed shape scores. Formally, for a batch of size *N* with predictions **y**^*_i_* and targets **y***_i_*, the loss function is:

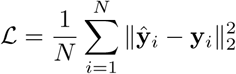

This loss was minimized using the AdamW optimizer [26] with a learning rate of 5 × 10*^−^*^6^ and weight decay of 10*^−^*^2^. Training was performed for up to 10,000 epochs using mini-batches of size 32. At each epoch, both training and validation losses were monitored, different hyperparameters were tested and the network state corresponding to the lowest validation loss was saved for evaluation.

#### Model Evaluation

Model performance was quantified using mean squared error (MSE) and the coefficient of determination (*R*^2^), computed per *def PC* component.

#### Geometric validation metrics

To quantitatively assess the geometric accuracy of the predicted deformations, reconstructed meshes from the predicted and true *def PC* scores were compared using several metrics:

- **Hausdorff distance:** The maximum distance from any point on the predicted mesh to its nearest neighbor on the true mesh, and vice versa. This measures the largest local discrepancy between two surfaces.
- **Average point-to-point distance (AvgDist):**The mean Euclidean distance from each point on the predicted mesh to the closest point on the true mesh. This quantifies the overall surface deviation.
- **Point-cloud overlap percentage (Overlap):** The proportion of points on the predicted mesh that lie within a 2 mm threshold of the nearest point on the true mesh. This metric indicates the fraction of the predicted surface that closely matches the true geometry.

Hausdorff distance, average point-to-point distance, and point-cloud overlap were computed using nearest-neighbor queries implemented via cKDTree from SciPy [27]. All metrics were computed separately for end-diastolic (ED) and end-systolic (ES) meshes and summarized as mean ± standard deviation across the test set. These measures complement volumetric comparisons and provide a detailed assessment of local and global geometric fidelity. To compare these measurements to the displacements of the models, the mean and maximum of the absolute displacements were recorded for each simulation run at both ED and ES with respect to the original shape. The average of both of these values was reported.

## 3 Results

### 3.1 Simulation Workflow

An example result of the FEM simulations is given in Fig 2. On the left is the volumetric mesh of the unloaded geometry. The mesh is then loaded with LVEDP = 1.20*kPa,* RVEDP = 0.533*kPa* to ED and then from ED to ES with LVESP = 18.0*kPa,* RVESP = 3.0*kPa* with a linearly ramped active tension set to 200.0*kPa*. The material properties were set to *a* = 1.28*kPa, a_f_* = 1.69*kPa*.

**Figure 2:**
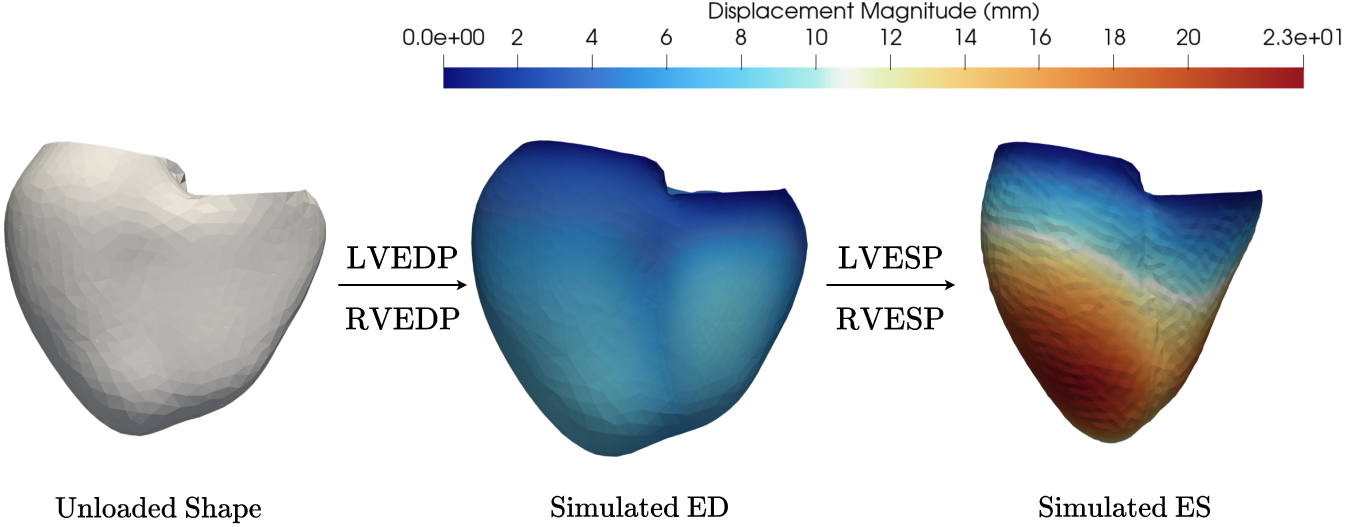
Simulation workflow from unloaded volumetric mesh to ED, and finally ES. Magnitude of the deformations are given in millimeter.

### 3.2 Hemodynamic Distributions

The physiological plausibility of the generated training data was assessed by comparing their density distribution with that of the metrics sampled from the UK Biobank. In Fig. 3, the simulated, sampled and UK Biobank population distributions of LVED and LVES volumes are compared showing reasonable overlap, the simulated ED volumes tended to have lower values than population values. Similarly, since we generated solutions for high ES pressures, ejection fractions skewed lower than the data but simulations covered all but the highest values above 65%.

**Figure 3:**
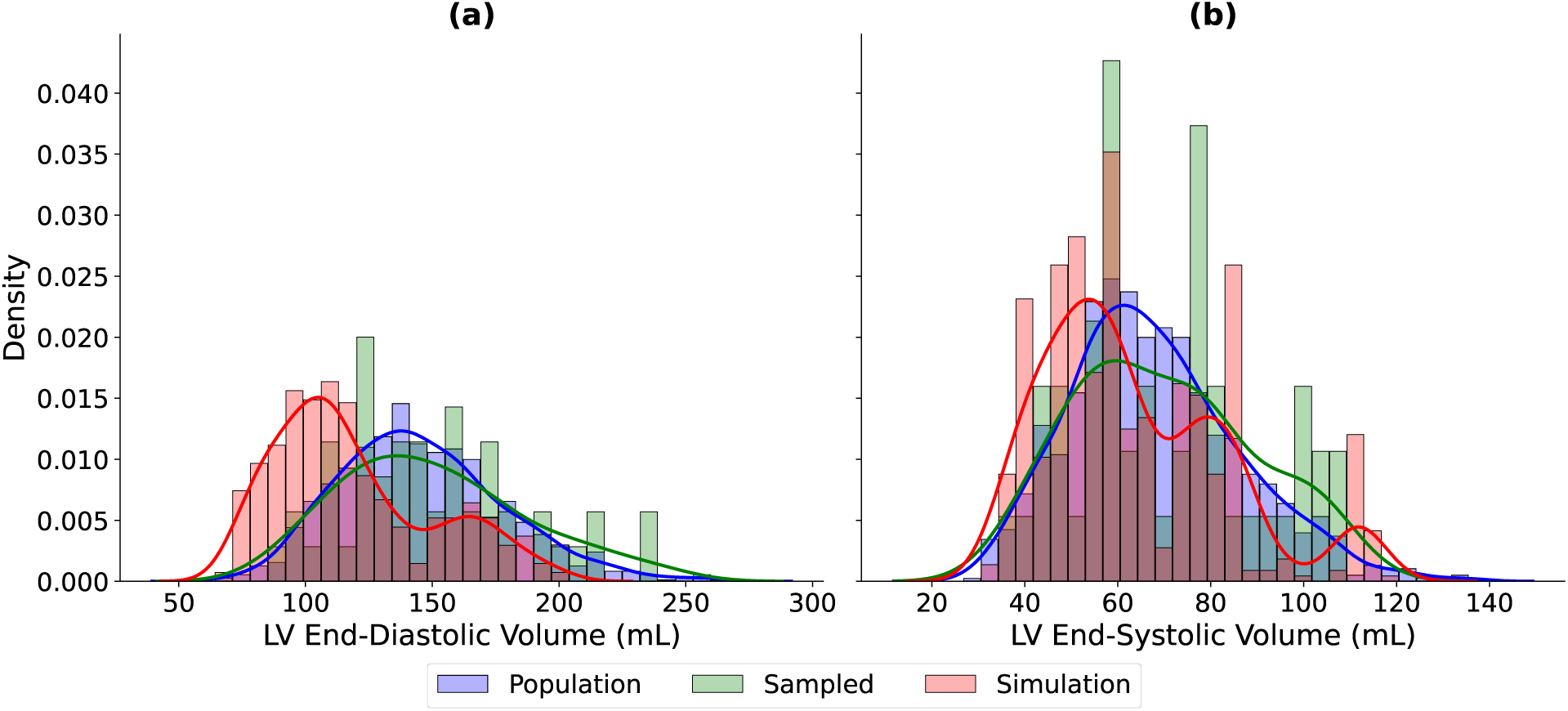
Comparison of left ventricular (LV) end-diastolic (ED) and end-systolic (ES) volume distributions between the population, sampled and simulation cohort. Histograms show the empirical volume distributions, while the solid lines represent smoothed kernel density estimates (KDEs) for the population (blue), sampled (green), and simulations (red). Panel (a) shows the distributions at ED, with means at 145.83, 150.35, and 118.49 mL for the population, sampled, and simulated cohorts, respectively. Panel (b) shows the distributions at ES with means at 67.95, 69.94, and 63.53 mL for the population, sampled, and simulated cohorts, respectively.

To better assess the function of the simulated hearts, the ejection fractions (EFs) of the population, sampled and simulated data were compared, as seen in Fig. 4a. The simulation results fall in the range of the atlas cohort. The EF of the simulation results were further analyzed by looking at the distribution broken down by prescribed LVESP. For the simulation, we used three conditions for LVESP as seen in Table 1, and these groups can be seen via Fig. 4b.

**Figure 4:**
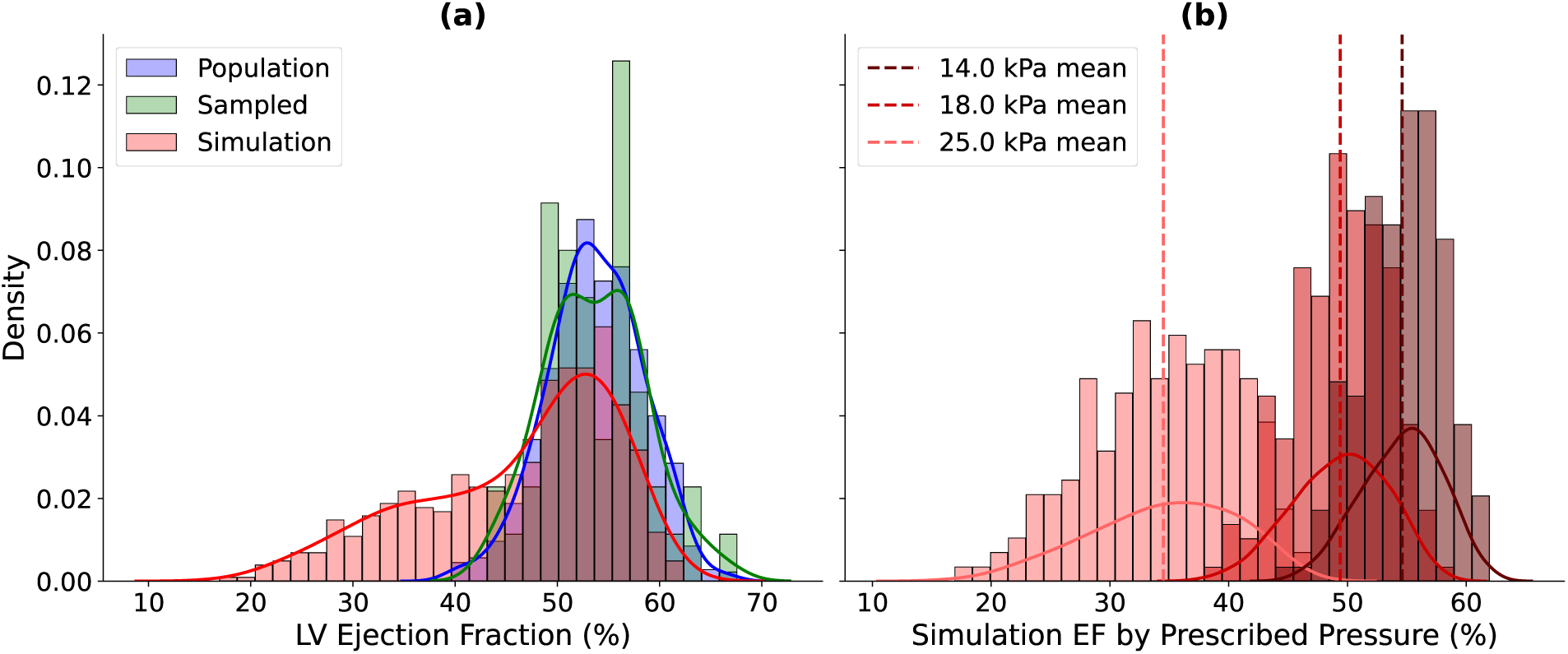
Comparison of left ventricular (LV) ejection fraction (EF) distributions. Histograms show the empirical EF distributions, while the solid lines represent smoothed kernel density estimates (KDEs) for the population (blue), sampled (green), and simulations (red). Panel (a) compares the three cohorts with means at 53.68%, 53.68% and 46.23% for the population, sampled and simulation cohort, respectively. Panel (b) shows simulation EF distributions grouped by prescribed end-systolic pressure (LVESP), with mean EF values marked as dashed vertical lines: 54.60% at 14 kPa, 49.40% at 18 kPa, and 34.50% at 25 kPa.

### 3.3 Training Set

The final count of simulations achieved can be seen in Tables 3 and 4 using the pressure conditions described in Table 1 and material properties (2.3), respectively. Overall, unloaded geometries corresponding to 50 different sampled cases were used to create the simulation data. The same unloaded geometries were repeated with different end-diastolic pressures and material properties to achieve a total 576 simulations. Additional unloaded geometries were incorporated into the training set by running select material parameter combination (e.g. *a* = 1.28*, a_f_* = 1.7) without having to cycle through all combinations. Since simulations with higher stiffness *a* converged more quickly, these material properties were somewhat over-represented in the final training set.

**Table 3:** Simulation counts by LV/RV pressure (kPa)

| PLVES | PLVED | PRVES | PRVED | Count |
| --- | --- | --- | --- | --- |
| 14 | 0.8 | 2 | 0.36 | 55 |
| 14 | 1.2 | 2 | 0.53 | 79 |
| 14 | 1.6 | 2 | 0.71 | 59 |
| 18 | 0.8 | 3 | 0.36 | 55 |
| 18 | 1.2 | 3 | 0.53 | 79 |
| 18 | 1.6 | 3 | 0.71 | 59 |
| 25 | 0.8 | 4 | 0.36 | 55 |
| 25 | 1.2 | 4 | 0.53 | 78 |
| 25 | 1.6 | 4 | 0.71 | 57 |
| Total Count |  |  |  | 576 |
| Total Geometries |  |  |  | 50 |

**Table 4:** Simulation counts by material properties (kPa)

| $a$ | $a_f$ | Count |
| --- | --- | --- |
| 0.50 | 1.68/1.69<sup>1</sup> | 80 |
| 0.50 | 10.0 | 69 |
| 0.50 | 20.0 | 60 |
| 1.28 | 1.00<sup>2</sup> | 3 |
| 1.28 | 1.68/1.69<sup>1</sup> | 185 |
| 1.28 | 10.0 | 89 |
| 1.28 | 20.0 | 90 |
| Total Count |  | 576 |

### 3.4 Surrogate Performance

For both the training and validation groups, mean squared errors decreased monotonically with epochs (Fig. 5). Validation loss stabilized right before 3000 epochs, and the model with the lowest validation loss was selected as the best-performing model and retained for evaluation.

**Figure 5:**
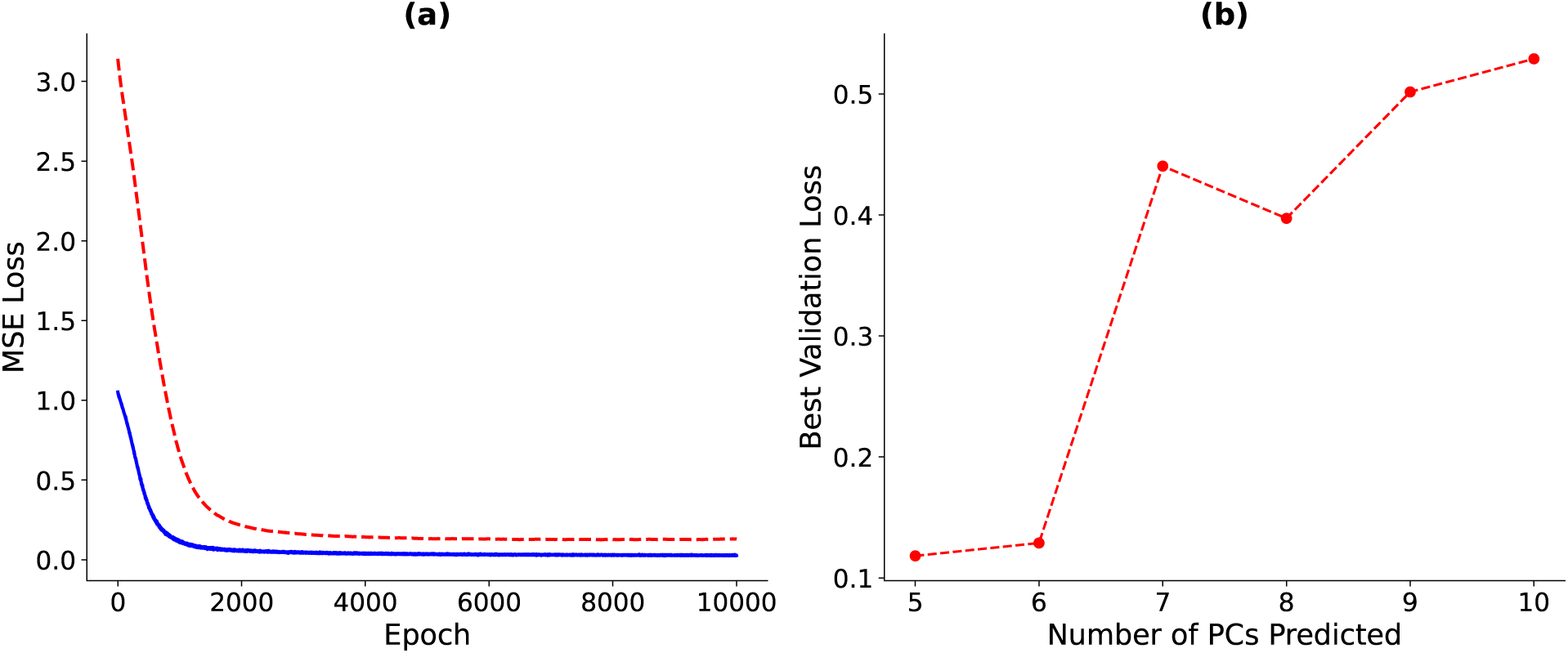
Training dynamics of the multilayer perceptron (MLP). (a) The figure shows mean squared error (MSE) loss across epochs for both the training (blue) and validation (red) datasets for predicting 5 PCs. (b) Best validation loss (MSE) as a function of the number of predicted principal components (PCs), shown in red.

Scatter plots of predicted versus true values were used to visualize the performance of MLP for each deformed principal component (*def PC*_1*−*5_) (Fig. 6). The scatter plots show most blue dots within their approximate 95% prediction intervals of the line of identity: ±0.49 (*def PC*_1_), ±1.40 (*def PC*_2_), ±0.41 (*def PC*_3_), ±0.42 (*def PC*_4_), and ±1.07 (*def PC*_5_). Discrepancies were greatest for *def PC*_2_.

**Figure 6:**
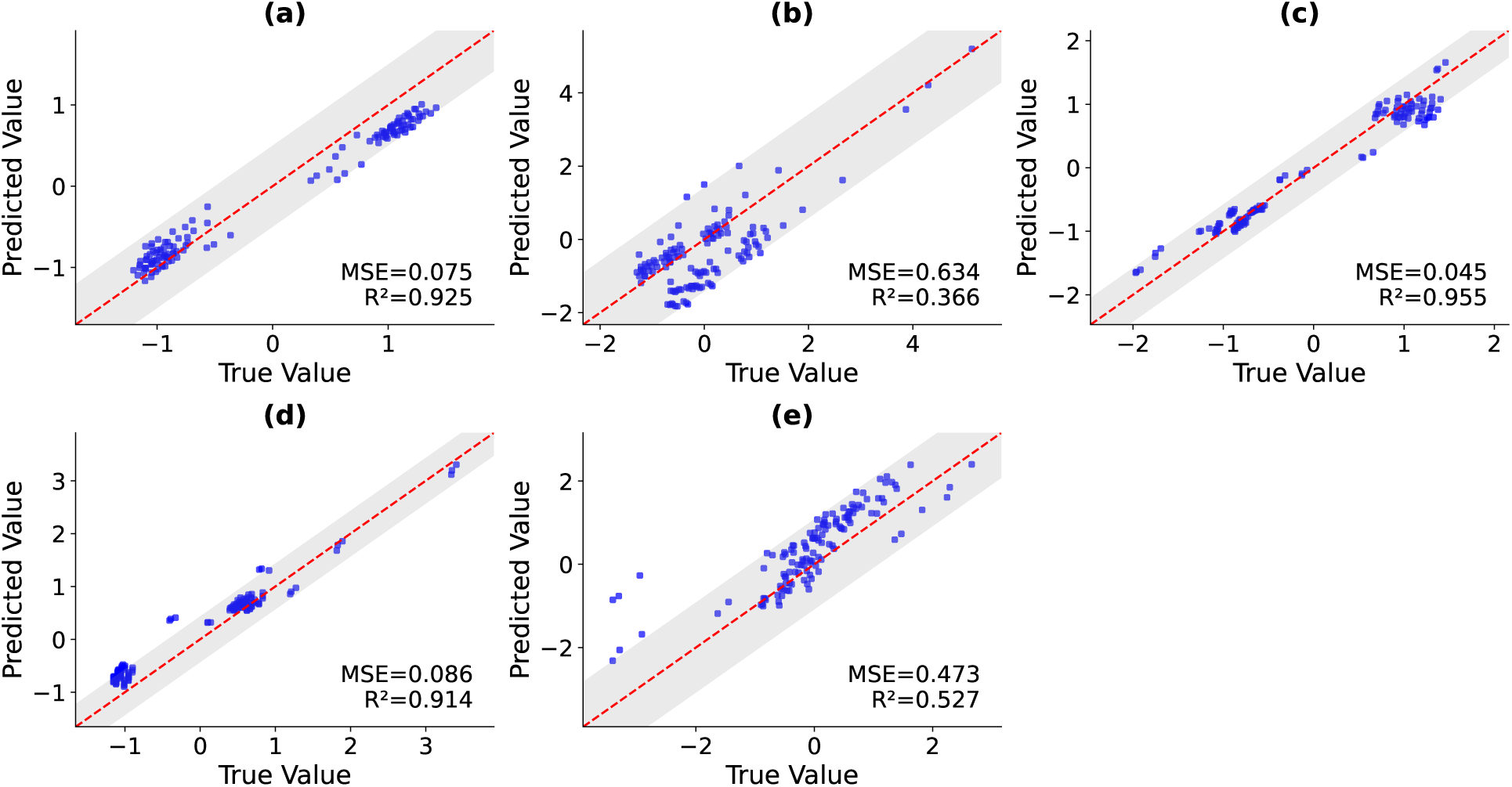
Scatter plots of predicted versus true values for each deformed PC (*def PC*_1*−*5_) in the test set. Panel a-e correspond to *def PC*_1*−*5_, respectively. Each plot shows predicted PCs on the y-axis vs. true values on the x-axis, with a red dashed line indicating the line of unity. The shaded gray region indicates the 95% prediction interval. The MSE and R² values for each component are reported in the plot.

The performance of the trained model was also validated using geometric comparisons with ground truth simulation data (2.6). RMS Geometric prediction errors were *<* 1 mm, worst-case errors (Hausdorff distance) were around 2 mm, and volumetric overlaps between predicted and ground truth deformed geometries were *>* 97% (Table 5). In comparison, the average mean and maximum absolute displacements of the training data from unloaded to ED and then to ES were 2.4 and 4.0 mm, and 10.7 and 18.2 mm, respectively.

**Table 5:** Geometric validation metrics of predicted vs true meshes. Values are reported as mean ± standard deviation. Distances are in millimeters (mm) and overlap in percent (%).

| Metric | Mean ± Std | Unit |
| --- | --- | --- |
| Hausdorff_ED | 1.69 ± 0.79 | mm |
| Hausdorff_ES | 1.81 ± 0.88 | mm |
| AvgDist_ED | 0.78 ± 0.37 | mm |
| AvgDist_ES | 0.77 ± 0.31 | mm |
| Overlap_ED | 97.58 ± 5.70 | % |
| Overlap_ES | 97.99 ± 4.18 | % |

## 4 Discussion

### 4.1 Key Findings

We found that a multilayer perceptron trained on PCA-based biventricular representations accurately predicts cardiac deformations, providing a fast and reliable surrogate for computationally expensive finite element simulations. This model was made possible by first developing a simulation workflow capable of producing physiologically plausible shapes, Fig. 2. As shown in Fig. 3, the simulated LV volumes fall within the range of the population. The mean simulated volumes, 119 mL and 64 mL, were slightly lower than the sampled population means of 146 mL and 68 mL for ED and ES, respectively. This difference may reflect the method used to compute the initial unloaded volumes (2.2) and the bias towards stiffer parameters in the training data (3.3). Exploring alternative approaches to defining the unloaded state or modifying material parameters could improve volume estimates; however, there are also many simulation constraints that can be fine-tuned. The effect of simulation inputs is also evident in ejection fraction (EF) distributions (Fig. 4). While the overall EF distribution spans the sampled population’s range of roughly 10–70%, it is not unimodal (Fig. 4a) like the population and sampled cohort. However, when grouping simulations by prescribed LVESP—14, 18, and 25 kPa(Fig. 4b)—the distributions within each cohort become unimodal, highlighting the strong dependence of EF on LVESP, with mean EF progressively decreasing as pressures rise, from 54.6% at 14 kPa to 49.4% at 18 kPa and 34.5% at 25 kPa, respectively. This trend falls in line with the expectation that a decrease in ES pressure will result in more ejection from the ventricle. This also suggests that producing additional simulations by linearly sweeping LVESP could improve robustness and better capture physiological variability.

However, it should be noted that the intention behind choosing the pressure and material parameters is not to match the training distributions of the population data, but to also span a wider range of pressures, volumes, and EFs. This broader coverage is necessary because the sampled data was based on data from volunteers who were mostly healthy.

Using 50 unloaded geometries sampled from the population distribution, the surrogate model was trained to predict deformed shapes from input parameters. Fig. 5a illustrates the minimization of mean squared error (MSE) for both training and validation datasets when predicting *def PC*_1*−*5_, which plateaued after approximately 3000 epochs, indicating convergence. Fig. 5b shows that as the number of *def PC*s included in the prediction increases, the best validation loss also increases even after 10,000 epochs. This trend suggests that the available dataset is insufficient to accurately predict higher-order PCs. While the validation loss remains below 0.2 for five and six *def PC*s, model performance is notably variable. A marked increase in loss occurs between predicting six and seven *def PC*s, indicating the onset of overfitting. Additionally, fluctuations in loss across predictions of *def PC*_1–8_ highlight the sensitivity of the small dataset to local minima. Overall the model showed good training and validation loss with predicting *def PC*_1*−*5_. Though insufficient for capturing the full extent of cardiac mechanics, it is a good start at understanding the range of data that will be needed to train a more extensive model.

When evaluating the ability of the model to predict *def PC*_1*−*5_ (Fig. 6), the model shows the highest correlation, R², with *def PC*_1_ at 0.94. This score corresponds to the size of the heart [6], and thus the model seems to be able to accurately predict the size of the heart. We see the worst correlation, R² of 0.451, with *def PC*_2_. This PC score is associated with apex-base height. The range for *def PC*_2_ in our simulated results was -5 to -1, which does not span the typical range of PC scores from about -3 to 3. This decreased correlation may be a result of the simulation parameters fixing the base’s displacement inadvertently affecting the *def PC*_2_ score. Because the atlas is built from real human data, it would be valuable to investigate the constraints placed on the bases in the simulations. Overall the mean R² of the model with all *def PC*_1*−*5_ was 0.7864 with a mean RMSE of 0.2114. Clearly this is skewed down by *def PC*_2_; however, this shows promising results that the model is performing well and with more training data it may improve.

The predicted and true *def PC*_1*−*5_ scores were also evaluated by translating the scores back into 3D geometries. This showed with sub-millimeter average surface errors compared to the average displacements of 2.4 and 10.7 mm at ED and ES and volumetric overlaps above 97%, demonstrating high fidelity to the simulated ground truth (Table 5).

### 4.2 Limitations

Despite promising overall performance, the surrogate model shows residual variability in certain PC modes, reflecting sensitivity to training data diversity and the limited size of the validation cohort. Due to the small training set, the model was trained only on the first 10 PC scores and predicts the first 5 PC scores, which explains only 78.2% and 63.5% of the total shape variability, respectively. Thus, its ability to reconstruct shapes corresponding to personalized data is currently truncated. These first 5 PC scores also may not be able to capture potentially important features of regional wall mechanics. The slightly lower simulated volumes compared to the sampled population suggest that both the method for calculating the unloaded state and the number of runs per condition may influence model fidelity. The unloaded shape was approximated using a simple linear heuristic. Although this may not significantly bias the surrogate model, subject-specific optimization will require a more accurate way of computing unloaded geometries consistent with the optimal material properties. This is an inverse dual problem itself. Additionally, reliance on prescribed LVESP for EF distributions indicates that the surrogate’s predictive accuracy is partially dependent on the range and density of simulation inputs. The simulations were also only optimized on two global passive material parameters (*a* and *a_f_*), whereas realistic optimizations would require spatially varying material properties, especially those corresponding to active contractile parameters.

Biases in the training data were also due to computational constraints. Simulations with stiffer material properties converged quickly and are therefore overrepresented, whereas less stiff material properties were excluded entirely due to the probative cost of smaller load steps. Thus, this resulted in a bias towards smaller geometries, stiffer hearts, and smaller EFs.

Finally, the boundary conditions applied on the tracts may have resulted in additional artifacts. The fixed Dirichlet boundary conditions (2.4) may be too restrictive, further resulting in smaller ED volumes and non-physiological deformations.

### 4.3 Future Work

Future work could expand the range of simulation conditions, including material properties, loading, and constraints, to increase the breadth of training simulations. The method of extracting the unloaded geometries could also be improved to be more consistent with the subject’s unique material properties. A more diverse and expanded training set would enable the development of models with additional unloaded and deformed PC scores, helping to improve the realism of the reconstructed models. It could also be trained as an inverse model and used to predict material properties from deformed PC scores.

## 5 Conclusion

These results demonstrate that machine learning surrogates can generate physiologically realistic cardiac deformations at a fraction of the computational cost of full finite element simulations. This approach promises to enable rapid subject-specific optimization and supports large-scale virtual studies, reducing the need for repeated full-scale simulations. This lays the foundation to using machine learning to solve the costly inverse challenge of determining material properties from biventricular end-diastolic and end-systolic geometries.

## Data Availability

Code is available upon request.

## Acknowledgments

We gratefully acknowledge the use of the cardiac atlas developed by Burns et al. [6]. This work was supported by the NIH grant NHLBI T32HL160507.

---

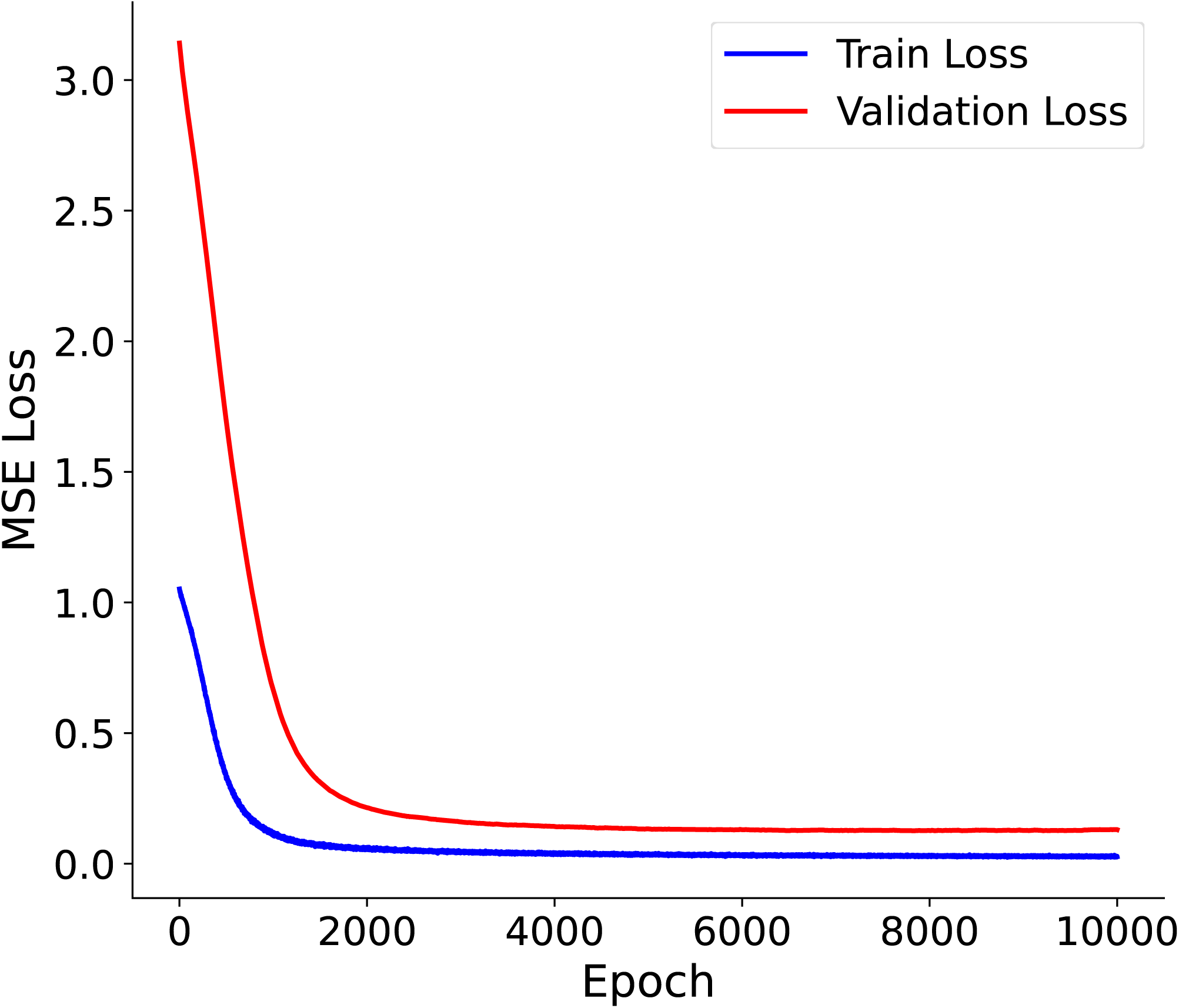

## Footnotes

^1^A combination of both values was included due to user error.

^2^This value was discontinued after one run due to issues with convergence, but was still included in the training set.

